# Geroprotective effects of *Aronia melanocarpa* fruit extract on *Drosophila melanogaster*

**DOI:** 10.1101/2021.03.03.433673

**Authors:** Elena Y. Platonova, Nadezhda V. Zemskaya, Mikhail V. Shaposhnikov, Denis A. Golubev, Daria V. Kukuman, Natalya R. Pakshina, Natalia S. Ulyasheva, Vasily V. Punegov, Sergey A. Patov, Alexey A. Moskalev

## Abstract

Aging and its consequences is one of the main problems of humanity. Phytochemicals such as polyphenols, flavonoids, anthocyanins have great potential as geroprotectors. Plants including Aronia used as dietary supplements are an excellent source of these compounds. We have studied the effects of the ethanol extract from Aronia fruits on *Drosophila melanogaster* lifespan, locomotor activity, stress resistance (oxidative stress, heat shock and starvation). And also, in order to reveal the influence of Aronia extract applied to different life cycles of *Drosophila melanogaster*, we selected different dietary schemes for imago: throughout life, during the first two weeks (1 – 2) and during average age (4 – 6 weeks). It was revealed that the ethanol extract of chokeberry increases the median life expectancy in males and females by 5% when *Drosophila melanogaster* is added to the diet for 4 – 6 weeks of life. This suggests that intervention even in old age is sufficient to increase lifespan. In addition, no harmful effects of ABE on locomotor activity were found as an indicator of fly health. We showed that Aronia extract increased stress resistance to hyperthermia and oxidative stress. At the same time, Aronia extract did not significantly affect the resistance of flies to starvation. ABE supplementation has increased expression of heat shock proteins (*Hsp27, Hsp68, Hsp83*), oxidative stress resistance genes (*Keap1, NRF, Sod1*), some circadian clock genes (*Clk, per*) and gene of longevity *Sirt1.*

## Introduction

Aging can be defined as a process that leads to a progressive decline in physiological integrity and organ dysfunctions (Van Beek et al. 2016). This decline is spreading at all levels of biological organization: molecular, cellular, tissue, and organismic (He and Jasper 2014a; Martel et al. 2019; Wang et al. 2014a). Hormesis is the stimulation of protective cellular mechanisms when exposed to mild stress (Rattan 2008) aiming in maintaining homeostasis that provides long-term positive effects (Calabrese et al. 2015; Rattan 1996; Rattan 2000). Hormesis inducers, so-called hormetins, can activate heat shock proteins and proteasomal degradation of damaged proteins (Beedholm et al. 2004; Verbeke et al. 2001a; Verbeke et al. 2001b) DNA repair (Martel et al. 2019) autophagy (Markaki and Tavernarakis 2011; Ryter and Choi 2013), an antioxidant defense that can promote lifespan increase (Pietsch et al. 2011; Rattan 2015). Hormetins may be food components, including phenolic acids, flavonoids, terpenoids, vitamins, trace elements which have a positive effect on the age-related quality of life (Calabrese and Blain 2005; Proshkina et al. 2020; Rattan 2015).

*Aronia melanocarpa* ((Michx.) Elliot) relates to the family *Rosaceae.* In height, the Aronia bush can reach up to 1 – 2 meters, its distinctive feature is dark red, almost black fruits in the form of an apple (Cvetanović et al. 2018; Hudec et al. 2006; Jurikova et al. 2017; Kokotkiewicz et al. 2010) In addition, black chokeberry is known all over the world due to its high content of various polyphenols, phenolic acids, flavonoids, anthocyanins, as well as vitamins and macro – and microelements (Borowska and Brzóska 2016; Cvetanović et al. 2018; Hudec et al. 2006; Jurikova et al. 2017; Kokotkiewicz et al. 2010; Oszmiański and Wojdylo 2005; Paulrayer et al. 2017; Staszowska-Karkut and Materska 2020; Taheri et al. 2013). In past centuries, both in North America and in Russia, natural products (berries, fruits, leaves, stems) were used in traditional medicine (Cvetanović et al. 2018; Jurikova et al. 2017; Kokotkiewicz et al. 2010). It was noted that the anti-inflammatory, antiviral and immunomodulatory effects of the use of decoctions, juices and infusions of chokeberry, which have a positive effect on the health and longevity of the organism (Jurikova et al. 2017; Kokotkiewicz et al. 2010; Staszowska-Karkut and Materska 2020). This means that society is interested in improving the quality of life and health with age, therefore, the current generation relies on the experience of the past, with the use of modern technologies.

The aging process is known to be associated with multiple changes in the body, including slow down of all biological processes, decrease of the reproductive activity and increase of inflammation (Flatt 2020; Rattan 2004). Recently, acetone extract of chokeberry fruits has shown to increase the median lifespan and locomotor activity in *Drosophila* (Jo and Imm 2017) The lifespan extending effect was associated with increased expression of superoxide dismutase (*SOD*), catalase (*CAT*), glutathione peroxidase (*GPx*) genes and suppression of the methuselah (*MTH*) gene (Jo and Imm 2017). It was shown that lifelong use of the polyphenol-rich extracts from blueberry (Peng et al. 2012), cloudberry (Lashmanova et al. 2019), rotten apple (Peng et al. 2011), cranberry (Wang et al. 2015) have positive effects on the *Drosophila melanogaster* lifespan. But the use of a geroprotector throughout life may be accompanied with increased risk of harmful side effects (Moskalev et al. 2016) and it is also known that the geroprotective effect persists when treated at a certain age. Therefore, in addition to experiments with the use of any substances throughout the life of model organisms, experiments were carried out where the effect of a substance on life expectancy was carried out only at an old age, in order to identify the greatest geroprotective potential of the studied substances. It was found that in adult mice (aged 20 – 22 months), when added to food, rapamycin exhibited the greatest geroprotective effect than in young mice (Wilkinson et al. 2012). The beneficial effects of chokeberry extracts have been revealed in different *in vitro* and *in vivo* models and at different levels from the cellular (Abdullah Thani et al. 2012; Cvetanović et al. 2018; Parzonko et al. 2015; Valdez et al. 2020) before organismic (Makanae et al. 2019).

The purpose of this work was to reveal the lifespan effects induced by ABE treatment at different ages. To assess whether the geroprotective effects of ABE can be affected by the age of treatment we supplemented Aronia extract to the *Drosophila* imago throughout life, during the ages of 1-2 weeks and 4-6 weeks of life, that correspond to young and average ages, respectively. We studied the effects of ABE treatment at different ages and concentrations on the healthspan (locomotor activity, intestinal integrity) and stress resistance. The maximum beneficial effect on lifespan was found for ABE treatment at the age of 4 – 6 weeks (3 – 9%). The performed RT-PCR assay revealed the effects of ABE on the age-related changes in the expression level of stress response genes. The ABE supplementation has increased expression of heat shock proteins (*Hsp27, Hsp68, Hsp83*), oxidative stress resistance genes (*Keap1, NRF, Sod1*), circadian clock genes (*Clk, per*) and gene of longevity *Sirt1* thereby increases resistance to oxidative stress and hyperthermia, but with age the expression of genes *Per*, *Sirt1* and *Keap1* has decreased. Therefore, ABE shows that different dietary schemes extended lifespan and resistance to different stress.

## Material and methods

### Extraction

Aronia berries were harvested in the autumn period (August - September), on the territory of the Komi Republic (Northwest Russia). The berries were pre-frozen at a temperature of − 20 °C. To prepare the extract, the fruits were crushed and centrifuged to obtain a supernatant. This mass was mixed with clay (0.1 molar hydrochloric acid solution) and centrifuged again. The resulting liquid was poured off and mixed with the extractant: 1% solution of concentrated hydrochloric acid in 96% ethanol. The resulting solution was centrifuged, and then ethanol from the extract was evaporated on an IR-1M vacuum rotary evaporator (Khimlaborpribor, Russia) at 35 ° C. The experimental concentrations of Aronia berry extract (ABE) were prepared from the obtained ethanol extract by dilution in 96% ethanol.

### High performance liquid chromatography (HPLC)

ABE samples were analyzed on a Thermo Finnigan liquid chromatograph (Thermo Fisher Scientific Inc., USA) using a diode array detector (200 – 600 nm) in tandem with a mass selective detector of the same company. The analysis was carried out at a wavelength of 520 nm, an eluent feed rate of 1 ml/min, analysis time of 40 minutes, in isocratic mode. The eluent acetonitrile - an aqueous solution of formic acid (10%) in the ratio of 7:93 (v/v) was used in the analysis). Column 4 × 250 mm with a sorbent Diasorb-130-C16T, granulation 7 μm.

For sample preparation 1 mg of the extract was dissolved in 10 ml of deionized water, after which it was applied to a prepared cartridge with the Hypersep C18 sorbent. The cartridge with the applied extract was washed with 10 ml of deionized water, then the target substances were washed off the cartridge with 1 ml of eluent (acetonitrile – aqueous formic acid solution (10%) in the ratio of 7:93 (v/v). The sample thus prepared was analyzed by High Performance Liquid Chromatography-Mass Spectrometry (HPLC-MS).

### *Drosophila* rearing

*D. melanogaster* wild type *Canton-S* line was obtained from Bloomington Stock Center at Indiana University (#64349, Bloomington, USA). The flies were maintained at 25 °C and at 60% relative humidity under a 12 h: 12 h light/dark cycle in a constant climate chamber Binder KBF720-ICH (Binder, Germany). The food media on which the flies lived contained water – 1000 ml, corn flour – 92 g, dry yeast – 32.1 g, agar-agar – 5.2 g, glucose – 136.9 g (Xia B et al. 2016) to which 5 ml of a 10% solution of nipagin in ethanol, and 5 ml of propionic acid were added.

### Treatment with Aronia berry extract

For imago feeding, ABE extracts (0.01; 0.1; 1.0; 2.5; 5.0 and 10 mg/ml) were directly added to the surface of the fresh medium (30 μl per vial). The 30 μl 96% ethanol was added to the medium surface of control vials. Vials were dried under a fan.

### Ages of ABE treatment

We treated flies with ABE at different imago ages. The first group received the extract throughout its life. The second group was treated at the age of 1-2 weeks old and the third group - at the age of 4-6 weeks old. The second and third groups, before and after the addition of the ABE, respectively, were fed with control medium. The experiments were carried out in 3 replicates.

### Lifespan analysis

After imago hatching, flies were anastezied using CO_2_, separated by sex, and transferred in vials containing the nutrient medium with investigated drugs (30 flies per vial) and the lifespan was assessed by recording the age of spontaneous death of flies. Dead flies were counted daily, and the remaining live flies were placed in new vials of fresh medium twice a week. The median and maximum (age 90% mortality) life expectancy and mortality doubling time (MRDT) were calculated.

### Stress resistance analysis

To investigate the effect of ABE feeding on the resistance to oxidative stress, starvation and hyperthermia, the newly enclosed male and female flies were collected and fed a diet with or without the ABE for 14 days and 33 days. The experimental flies were reared on the nutrient medium with ABE in concentrations 0.1, 1.0 and 5.0 mg/ml. To assay resistance to oxidative stress, flies were exposed to medium composed of 2% agar, 5% sucrose and 20 mM paraquat (Sigma-Aldrich, USA). During starvation the flies were kept on 2% agar medium. Hyperthermia was induced by continuous exposure of the flies to 35°C. Dead flies were identified using the DAM2 *Drosophila* Activity Monitor (TriKinetics Inc., USA) by the complete absence of movement. In each experimental variant 32 flies of each sex were analyzed. All experiments were carried out in two replicates.

### Analysis of locomotor activity

The age-dependent changes in spontaneous locomotor activity was measured using the LAM25 Locomotor Activity Monitor (TriKinetics Inc., USA) under standard 12 h lights on, 12 h lights off conditions. The data from 10 flies in 4 vials as replicates were collected during 24 h and represented as average total daily locomotor activity. Measurements were carried out every week, from the age of 1 to 9 weeks. The experimental flies were contained on the nutrient medium with ABE in concentrations 0.1, 1.0 and 5.0 mg/ml.

### Analysis of intestinal integrity

The Smurf assay was used for intestinal barrier permeability estimation (Rera et al. 2012). The flies from both control and experimental cohorts were tested at ages of 2, 6 and 8 weeks. Both cohorts were kept during 16 hours on a food medium containing 2.5% (mass/volume) blue food dye (Brilliant Blue FCF) after that the flies were transferred to the standard medium. The induction of “Smurf”-phenotype detecting intestinal permeability increase was accounted for in the sample if only the whole fly was dyed blue. The experimental flies were contained on the nutrient medium with ABE in concentrations 0.01; 0.1; 1.0; 2.5; 5.0 and 10 mg/ml.

### Analysis of Food choice (FLIC assay)

Time-dependent changes in food preference were analyzed using flies at the age of 7 and 45 day old from the control and ABE-treated groups. A fly liquid – food interaction counter (FLIC) system (Sable Systems, USA) was used to analyze feeding behavior as described (Ro et al. 2014). For food choice assay each channel of *Drosophila* Feeding Monitor (DFM) was loaded with either 5% sucrose (−ABE control) or 30 μl/ml ABE in 5% sucrose (+ABE experiment). Before the assay, flies were starved in *Drosophila* vial without medium during 2h 30m. The assay was performed for 3 hr using 6 flies. FLIC Monitor Software (downloaded from flidea.tech) was used to collect raw data from the DFM. Feeding Preference Index (PI) values from the FLIC system were calculated as the difference in total feeding time between the control and experimental foods divided by total feeding time for both foods (Ro et al. 2014). The PI ranged from 1 (complete preference for −ABE control food) to −1 (complete preference for +ABE experimental food) with a value of 0 indicating no food preference. PI value was calculated for each individual fly and presented as mean PIs for the experimental groups.

### Quantitative real time PCR

For each variant of the experiment, 30 males and females were selected per test tube (Genesee Scientific, USA) on the nutrient medium with ABE in concentrations 0.1, 1 and 5 mg/ml. Males and females lived separately at 25 °C under 12-hour lighting in Binder KBF720-ICH climatic chambers (Binder, Germany). The analysis was performed at the age of 14 and 33 days. For each variant of the experiment, 20 males and 10 females were selected. The experiment was carried out in three biological replicates, with three analytical replicates inside each.

Gene expression was measured by quantitative real-time PCR with a reverse transcription step. RNA was isolated using an Aurum Total RNA mini kit (Bio-Rad, USA) according to the manufacturer’s instructions. RNA concentration was measured using a Quant-iT RNA Assay Kit (Invitrogen, USA) according to the manufacturer’s instructions. cDNA was synthesized according to the iScript cDNA Synthesis Kit (Bio-Rad, USA) from the resulting RNA solution. The reaction mixture for the PCR reaction was prepared according to the instructions for iTaq Universal SYBR Green Supermix (Bio-Rad, USA) and primers (Supplementary Table S5). The primer design was performed using QuantPrime online tool (Arvidsson et al. 2008). The polymerase chain reaction was carried out using primers (Supplementary Table S5) in a CFX96 amplifier (Bio-Rad, USA) using the following program: 1) 95 ° C for 30 s, 2) 95 ° C for 10 s, 3) 60 ° C for 30 s, 4) steps 2-3 were repeated 49 times 5) DNA melting step.

The expression of the studied genes was calculated relative to the expression of the housekeeping genes Tubulin, eEF1α2, RpL32 using the CFX Manager 3.1 software (Bio-Rad, USA). Differences were considered significant changes (increase or decrease) in expression: p <0.05 according to the Student’s t-test and crossing the Regulation Threshold.

### Statistical analysis

To compare the statistical differences in survival functions and median lifespan between control and experimental groups, the modified Kolmogorov-Smirnov and log-rank test were used, respectively (Fleming et al. 1980; Mantel 1966). A Wang-Allison test was used to estimate the differences in the age of 90% mortality (Wang et al. 2004a). To assess the statistical significance of differences in resistance to stress factors, the Fisher’s exact test was used (Mehta et al. 1984). A Mann Whitney U-test was used for pairwise comparisons of feeding preference indexes (PMID: 11509435). Statistical analyses of the data were performed using STATISTICA software, version 6.1 (StatSoft, USA), R, version 2.15.1 (The R Foundation) and OASIS 2 (Online Application for Survival Analysis 2) (Han et al. 2016). R code for simple FLIC analyses Version 4.0 was used to analyze the signal data from FLIC DFMs (https://github.com/PletcherLab/FLIC_R_Code) (Ro et al. 2014).

## Results

### Chemical analysis of berry extracts

To determine the active substances in the ethanol extract of chokeberry, an HPLC method was required, which was carried out at a wavelength of 520 nm, an eluent flow rate of 1 ml/min and an analysis time of 40 minutes in an isocratic mode (Fig 1).

**Fig. 1.**
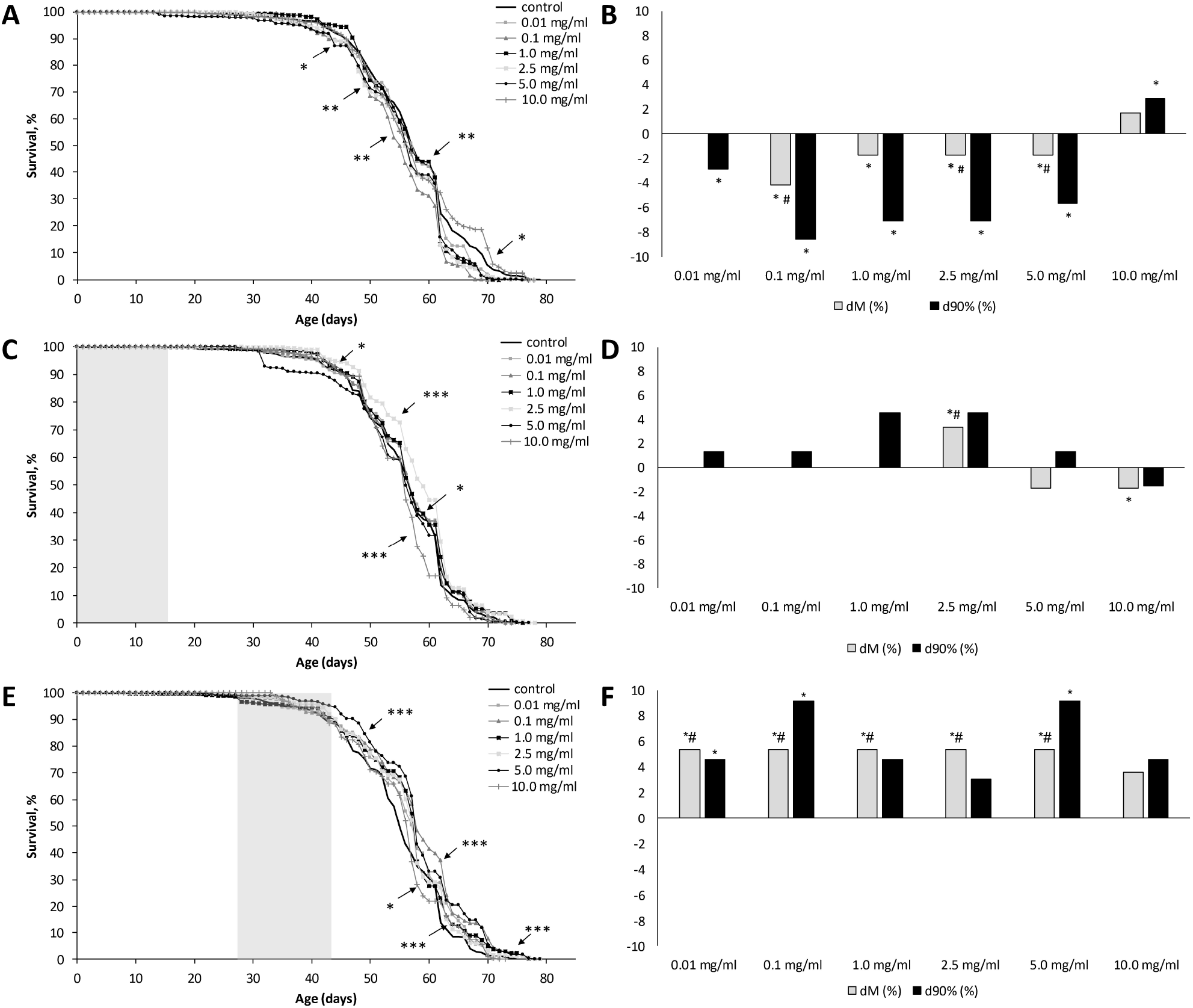
High performance liquid chromatography sample “Aronia”: (1) 10.54 min - delphinidin (Glu); (2) 13.38 min - delphinidin (Rut); (3) 16.64 min - cyanidin (Glu)

The chromatogram (Fig 1) shows the main jumps: at 10.54 min - delphinidin (Glu), 13.38 min - delphinidin (Rut), 16.64 min - cyanidin (Glu), which indicates that these substances are the main ones in the analyzed extract. Delphinidin and cyanidin are anthocyanidins, they are found in many bright blue-red fruits and berries, such as Manitoba berries, Saskatoon berry (Hosseinian and Beta 2007), eggplant (Sigurdson et al. 2018), more delphinidin will be collected. Cyanidin is the most common in raspberries, strawberries (Hosseinian and Beta 2007), and black chokeberry (Jurikova et al. 2017), sour cherry (Ertan et al. 2018) in apple skin (Ban et al. 2009), acai (Yamaguchi et al. 2015).

Unlike other fruits and berries, aronia fruits can contain a wide range of biologically active compounds, such as polyphenols: phenolic acids, flavonoids, anthocyanins, proanthocyanidins in various proportions (Tolić et al. 2015). This largely depends on the time of collection, on soil factors, climatic conditions, as well as on the extraction method (water, acetone, ethanol, water-ethanol) (Cvetanović et al. 2018; Hudec et al. 2006; Jurikova et al. 2017; Kokotkiewicz et al. 2010)

The effects of treatment with Aronia berry extract (ABE) at different ages of the imago The aging is associated with multiple physiological disorders. While reproductive potential and locomotor activity decrease, the inflammatory reactions and pathological processes increase (Flatt 2020; He and Jasper 2014b). Thereby the effect of geroprotective intervention may depend on the age of the organism. To reveal the connection between the geroprotective effect and age we studied the lifespan effects of ABE when imago flies were treated at different ages. In addition, it is necessary to take into account the characteristics of the periodization of the fly’s life cycle, during which the most important changes in the body occur: reproductive maturation (the first days of life), then reproductive activity, then a decrease in reproductive activity and further death (Flatt 2020). Therefore, we chose different treatment ages, namely: (1) throughout life, (2) at the age of 1 – 2 and (3) 4 – 6 weeks at different concentrations (0.01; 0, 1; 1.0; 2.5; 5.0 and 10 mg/ml).

It was revealed that the ethanol extract from the fruits of chokeberry in concentrations: 0.01; 0.1; 1.0; 2.5; 5.0 and 10.0 mg/ml, supplied with food throughout life, negatively affects the median and the maximum lifespan (LS) of male (see Fig. 2A, B) and female (see Fig. 3A, B) *Drosophila melanogaster* (Supplementary Table S1). When the extract was added to food for 1 – 2 weeks of adult life, 2.5 mg/ml in males (Fig. 2C, D) slightly increased the median lifespan (by 3%), and in females (Fig. 3C, D) 10.0 mg/ml reduced the median LS of fruit flies by 9% (Supplementary Table S2). Adding the extract to food at 4-6 weeks of adult life in males (Fig. 2E, F) increased the median lifespan by 5%, and the maximum LS was increased by 9% at concentrations of 0.1 and 5.0 mg/ml. In females (Fig. 3E, F), 0.1 and 5.0 mg/ml prolonged the median lifespan by 5%, and all concentrations except 10.0 mg/ml increased the maximum lifespan of fruit flies by 3% (Supplementary Table S3). Based on the data obtained, the use of the extract in middle age (4 – 6 weeks of life) has the greatest effect in increasing the lifespan in *Drosophila melanogaster* than its use throughout life and in the first weeks of life.

**Fig. 2.**
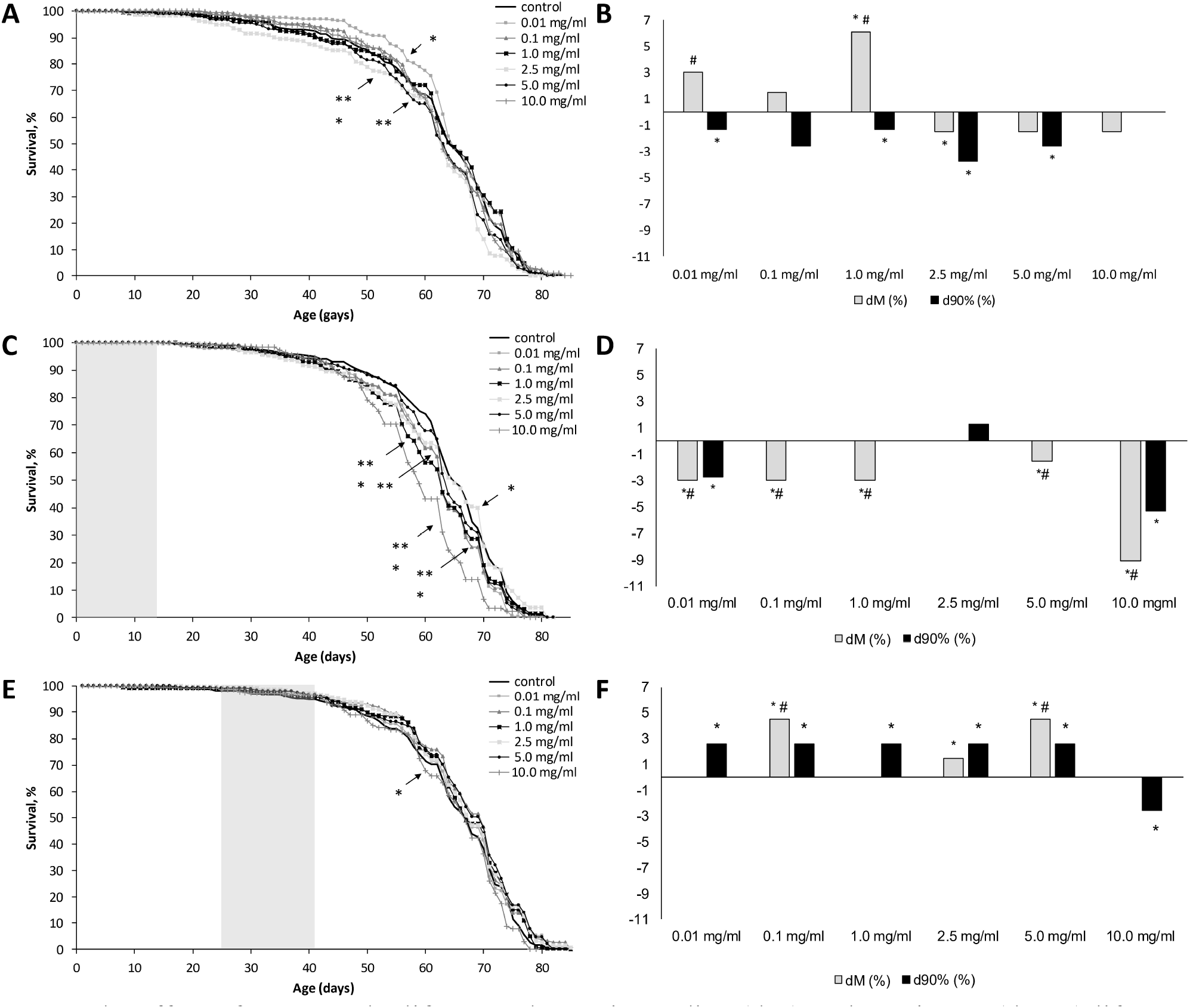
The effect of ABE on the lifespan, change in median (dM) and maximum (d90%) lifespan in males of *D. melanogaster*, throughout the whole lifetime of imago (A, B); at the age of 1-2 weeks old (C, D) and at the age of 4-6 weeks old (E, F). The results of the three replications are presented. The gray background shows the ages of ABE treatment. * p<0.05 - Kolmogorov-Smirnov test; *p<0.05 - Mantel-Cox test for median lifespan; Wang-Allison test - maximum lifespan; #p<0.05 - Wilcoxon-Breslow-Gehan test for median lifespan

**Fig. 3.**
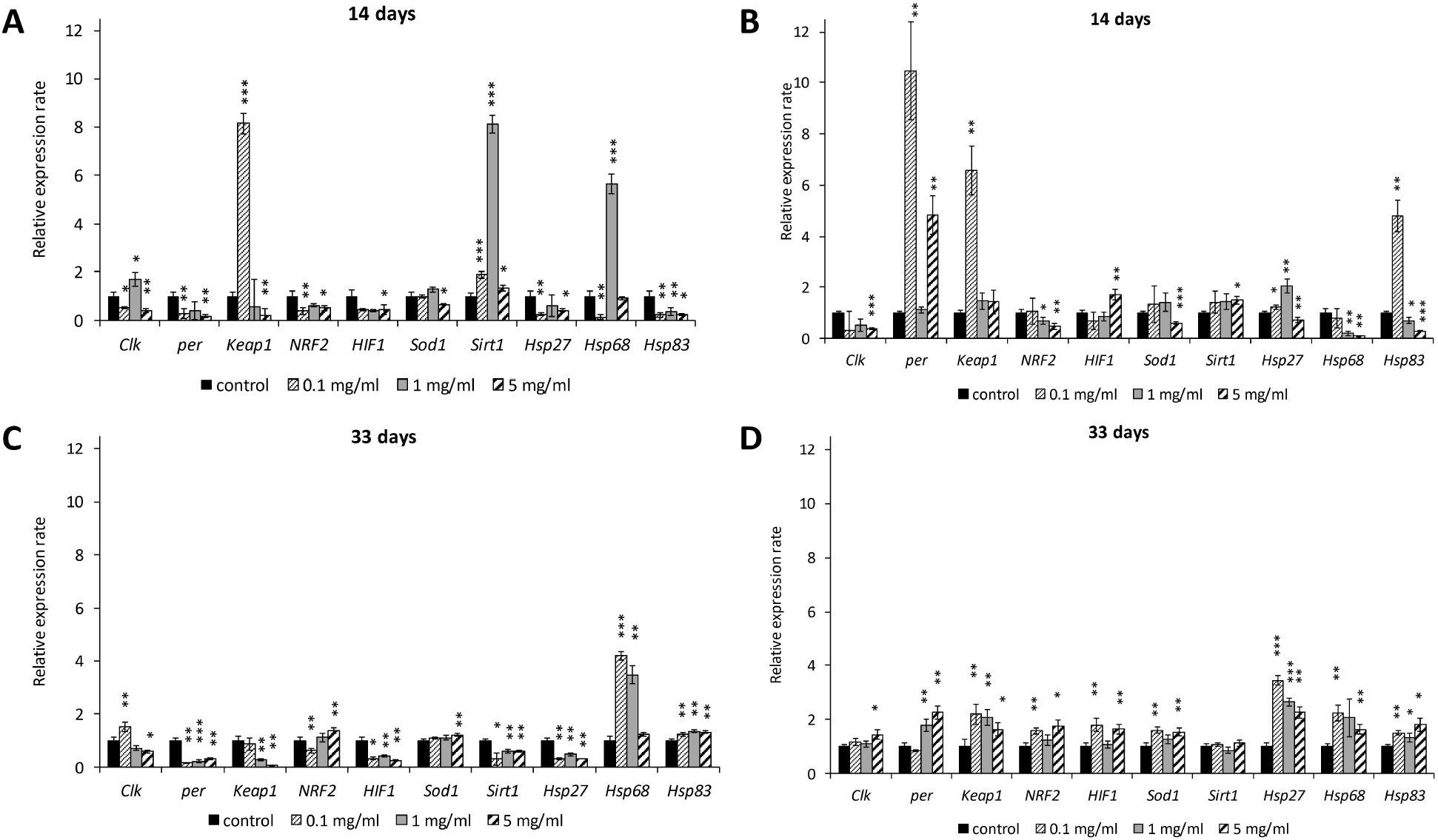
The effect of ABE on the lifespan, change in median (dM) and maximum (d90%) lifespan in females of *D. melanogaster*, throughout the whole lifetime of imago (A, B); at the age of 1-2 weeks old (C, D) and at the age of 4-6 weeks old (E, F). The results of the three replications are presented. The gray background shows the ages of ABE treatment. * p<0.05 - Kolmogorov-Smirnov test; *p<0.05 - Mantel-Cox test for median lifespan; Wang-Allison test - maximum lifespan; #p<0.05 - Wilcoxon-Breslow-Gehan test for median lifespan

Previously, the effect of berry extracts used throughout the life of an imago on the model organism of *Drosophila melanogaster* was studied. Powder extract from bilberry (2.0 and 5.0 mg/ml) increased the average lifespan by 10% in *Drosophila melanogaster* (Peng et al. 2012) Acetone extract of cloudberries (0.12 mg/ml) in females caused an increase in median lifespan by 11% (Lashmanova et al. 2019). Polyphenol-rich apples increased average lifespan by 10% (Peng et al. 2011). Anthocyanin extract of cranberry (20 mg/ml) also increased the average lifespan of males by 10% (Wang et al. 2015). In an experiment with a mouse model with various age-related pathologies, individuals at the age of 20 – 22 months received doses of rapamycin with food. In the group of mice receiving rapamycin, it showed a geroprotective effect, namely, reduced the risk of developing age-related diseases (changes in the liver, heart, adrenal glands) compared with the control group (Wilkinson et al. 2012)

In our experiments, we confirmed that to increase the lifespan of *Drosophila melanogaster*, a short intervention is enough even in old age. And when the extract is used throughout life, it has a negative effect, perhaps this is due to the extraction method and its contents.

#### Locomotor activity

Aging is associated with a decrease in the functionality of the whole organism and in particular the locomotor function (Wang et al. 2014b). Therefore, to reveal the potential efficiency of chokeberry extract to delay the functional aging of flies, we investigated the effect of the most effective geroprotective concentrations (0.1; 1.0 and 5 mg/ml) on the age-related changes of locomotor activity in *Drosophila melanogaster*.

No statistically significant differences in locomotor activity under the influence of chokeberry extract were revealed in males (Fig. 4A). However, in females (see Fig. 4B) 5 mg/ml ABE was the most effective concentration, which increased their locomotor activity at the age of 4 weeks old. At the same time, no harmful effects of ABE on locomotor activity as an indicator of the health of flies were found.

**Fig. 4.**
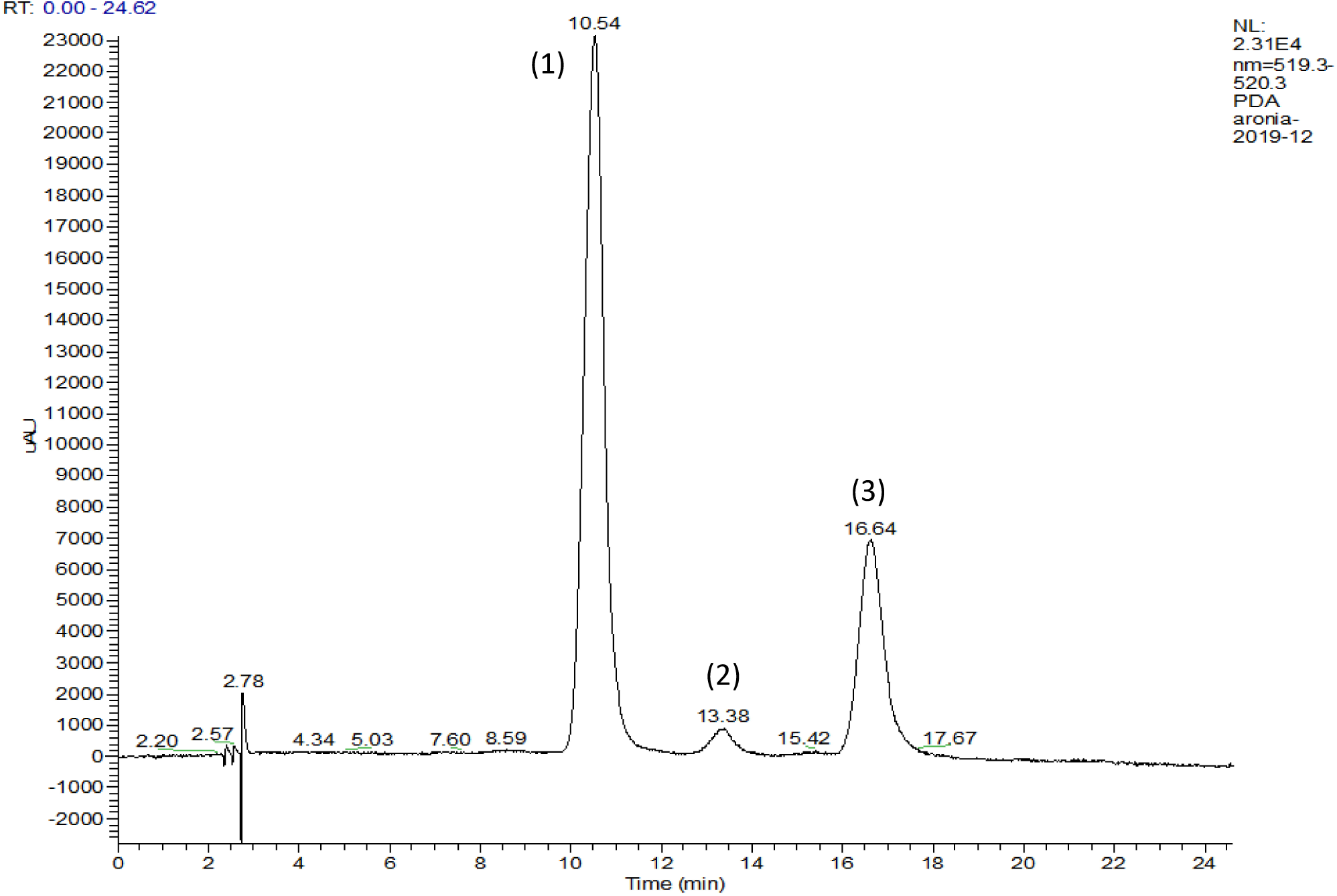
Effects of the ABE on the age-related changes of locomotor activity in *Drosophila* males (A) and females (B), *p<0.05, Student’s t-test.

One of the previous studies showed that Aronia supplement at a concentration of 2.5 mg/ml also useful for maintaining locomotor activity in male flies (Jo and Imm 2017). This suggests that Aronia supplement may increase not only lifespan, but healthspan also. However, in several other works, berry supplements had no effect on locomotor activity in rats (Eftimov and Valcheva-Kuzmanova 2018; Fernández-Demeneghi et al. 2019; Janšakova et al. 2016; Valcheva-Kuzmanova et al. 2014; Valcheva-Kuzmanova and Zhelyazkova-Savova 2009) (Pomatto et al. 2017). So the question has been raised, what exactly is the mechanism of influence on locomotor activity by berries? The posed question needs to be verified by other methods.

#### The Smurf assay

Under the influence of external (changes in diet and lifestyle) and internal (various diseases) factors, the permeability of the intestinal barrier decreases (an increase in the number of inflammatory reactions and a decrease in the protective function) which negatively affects the health and quality of life of the body, especially with age. That is, by improving the functions of the intestinal barrier, namely its permeability, it is possible to reduce the number of age-dependent diseases and increase lifespan (Bischoff et al. 2014; Rera et al. 2012; Vancamelbeke and Vermeire 2017). We have studied the effect of ABE on the *Drosophila* intestinal barrier permeability at the age of 2, 6 and 8 weeks old (Supplementary Table S4). At the age of 2 weeks, flies with the “Smurf” phenotype were not found. But at the age of 6 weeks flies and 8 weeks with the “Smurf” phenotype were identified. This suggests that with age, the functioning of a living organization decreases, including a violation of the integrity of the intestine, which in the future can lead to the death of the organism (Rera et al. 2012). But, no significant changes in males and females were found at 2, 6 and 8 weeks of life, which suggests that the ethanol extract of chokeberry with concentrations of 0.01; 0.1; 1.0; 2.5; 5.0 and 10.0 mg/ml had no negative effect on the permeability of the intestinal barrier. Previously the effect of various chokeberry extracts on gastric permeability has been studied. Valdez et al, shows that aronia polyphenolic powder improved (0.5–10 mg/ml) barrier function of inflamed Caco-2 cells (Valdez et al. 2020). Olejnik et al, *in vitro* experiments on Caco-2 cells, using the *Sambucus nigra L*. fruit extract passing through the gastric tract, had an anti-inflammatory effect (Olejnik et al. 2016). Paulrayer et al found a gastroprotective effect in rats at a concentration of 200 mg/kg of an aqueous alcoholic extract of chokeberry (Paulrayer et al. 2017). From the above studies, it can be assumed that chokeberry extracts have a protective effect on the gastrointestinal tract, which is directly related to the quality of health, which ultimately can reduce age-related diseases (chronic inflammation, metabolic syndrome, insulin resistance) (Rera et al. 2012). Further study is needed of the effects of chokeberry extract, with a possible decrease in concentration, on the permeability of the intestinal barrier in *Drosophila melanogaster*.

#### Food choice (FLIC assay)

The high levels of phenolic compounds in Aronia may contribute to pronounced bitterness, sourness and astringency of ABE containing food media (Duffy et al. 2016; Sidor and Gramza-Michałowska 2019) and influence the feeding rate (Ro et al. 2014). Both reduced or excessive food consumption may affect nutritional status, health, and finally the lifespan (Tatar et al. 2014; Wong et al. 2009). To find out how ABE supplementation affects the food consumption we studied dietary preferences of flies at different ages using the FLIC system (Ro et al. 2014). In addition we revealed the possible effect of habituation to chokeberry flavor by analyzing feeding behaviour in flies that were kept either on control media or on media with ABE before the test.

The obtained results reflect the lack of strong preference (mean PI varies in wide ranges) of the males (0.01 mg/ml ABE) and females (0.01, 2.5, and 10 mg/ml ABE) for −ABE or +ABE food medium (Fig. 5). This data confirms the results of a previous study that demonstrated no significant difference in food intake between flies from control and supplemented with 2.5 mg/ml aronia acetone extract groups estimated by gustatory assay (Jo and Imm 2017) demonstrating that the observed effects on lifespan and on healthspan are not associated with changes in food consumption. At the same time males of all groups exhibited weak preference (mean PI~0.2) in favor of control (−ABE) medium over 10 mg/ml ABE (Fig. 5E), suggesting that at high concentrations, the intense taste of chokeberry extract can lead to reduced food intake and induce dietary restriction-mediated effects. Control and ABE pretreated 7 days old males exhibited statistically significant (p<0.05) weak preference (mean PI~−0.3) in favor of 2.5 mg/ml ABE media compared to 45 days old males (Fig. 5C). These age-related changes in food preference may partly explain the differences in lifespan effects of ABE related to feeding at different ages.

**Fig. 5.**
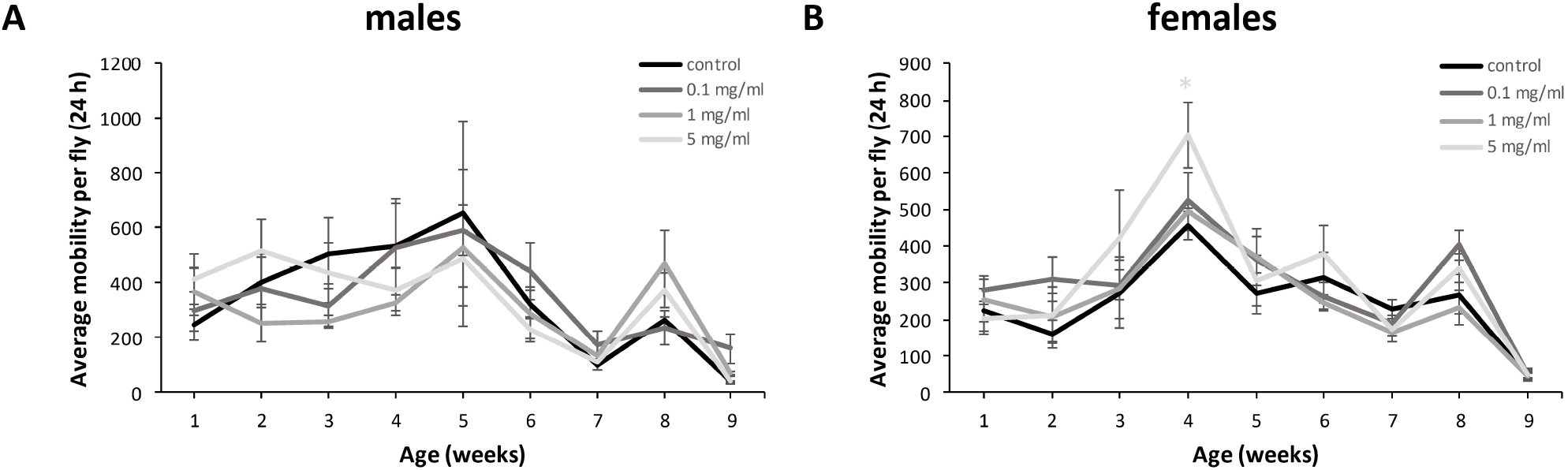
The effects of ABE supplementation on the food choice estimated by feeding Preference Index (PI) in males (A, C, E) and females (B, D, F). ABE was supplemented in liquid food obtained by dilution of 30 µl of 0.01 mg/ml (A, B), 2.5 mg/ml (C, D), and 10 mg/ml (E, F) ABE in 1 ml of 5% sucrose solution. PI = 1 – complete preference for –ABE control food; PI = −1 – preference for +ABE experimental food; PI = 0 indicating no food preference. *p<0.05, Mann Whitney U-test

#### Stress resistance

The mechanisms of resistance to stress and longevity are interrelated (Saunders and Verdin 2009). Oxidative stress, which leads to disruption of redox homeostasis, is one of the important factors of aging and the organism’s ability to respond to oxidative stress plays an important role in longevity (Kuether and Arking 1999). Treatment of flies with paraquat, high temperature and starvation may be a cause of free radicals, which in the future can lead to pathological processes in the body and further death (Nordquist et al. 1995; Robinson et al. 1997; Turrens 2003). In addition decreased stress tolerance is one of the manifestations of aging (Ikeyama et al. 2002; Pandolf 1997). Therefore, we were interested in how the aronia extract will affect the resistance of *Drosophila melanogaster* to various stresses (oxidative, hyperthermia and starvation) at the age of 14 and 33 days old (see Fig. 6). In these assays we used the extract at concentrations of 0.1; 1.0 and 5.0 mg/ml, that demonstrated the most effective geroprotective potential.

**Fig. 6.**
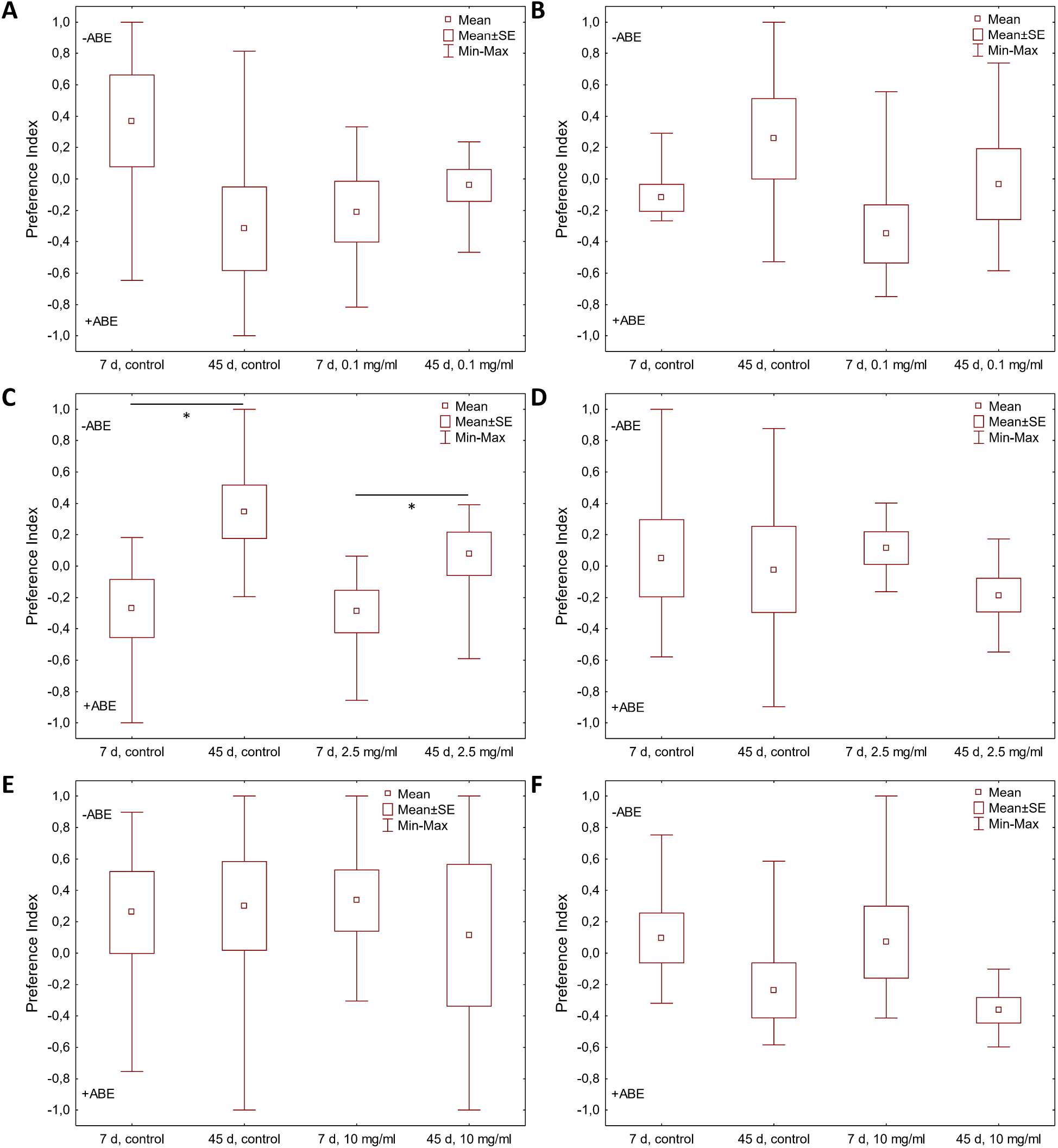
Effects of the ethanolic extract from berries of *Aronia melanocarpa* on the resistance of *Drosophila* males (A, C, E) and females (B, D, F) at the age of 14 and 33 days old to oxidative stress (A, B), starvation (C, D), and hyperthermia (E, F). *p<0.05 Fisher’s exact test

According to our data, in males (Fig. 6A), 1.0 mg/ml ABE increased resistance to oxidative stress at 33 days of life but significantly reduced resistance to starvation at 14 days (Fig. 6C). And 0.1 and 5.0 mg/ml ABE significantly increased the resistance of males to hyperthermia (Fig. 6E) on the 14th day of imago life. In females, 5 mg/ml ABE had a protective effect against oxidative stress at 33 days of age (Fig. 6B), as well as starvation at 14 days of age in flies (Fig. 6D). On the 14th day of females’ life, 0.1 and 5.0 mg/ml ABE increased resistance to hyperthermia, but on the 33rd day, all concentrations had a negative effect on resistance to hyperthermia (Fig. 6F). It is well known, *Keap1* suppresses transcription activity of *Nrf2* (Jaramillo and Zhang 2013). Keap1/NRF2 signaling protects the body from oxidative stress, including aging-related diseases (Cheng et al. 2016; Li et al. 2019; Martel et al. 2019; Sykiotis and Bohmann 2008). We have shown an increase the resistance to oxidative stress of *Drosophila* at 14 and 33 days after treatment with ABE, perhaps it is due to the activation of *Nrf2* and the subsequent decrease in the level of expression of *Keap1*. At the same time, Aronia extract did not significantly affect the resistance of flies to starvation, a similar effect was shown on cloudberries (Lashmanova et al. 2018).

Findings on the effect of ABE on the resistance to hyperthermia, it can be concluded that when Aronia extract is obtained for 14 days, both males and females, stress resistance increases, but already on day 33 their stress resistance decreases in both males and females. Perhaps due to prolonged and/or intake of high doses, flavonoids begin to act as mutagens, prooxidants, thereby aggravating their quality of life (Skibola and Smith 2000).

##### RT-PCR

Changes in gene expression levels are associated with the health and lifespan (Tyshkovskiy et al. 2019). Plant extracts have a wide range of pharmacological effects, which at the molecular level are determined by the effects on the transcriptional activity of genes, including stress response genes. Thus, we have studied the changes in expression level of cellular stress response genes, including heat shock proteins (*Hsp27, Hsp68, Hsp83*), oxidative stress resistance genes (*Keap1, NRF, Sod1, HIF1*) and circadian clock genes (*Clk, per*) gene of longevity *Sirt1*. The mRNA expression level was identified by RT-PCR on 14 (Supplementary Table S6) and 33 (Supplementary Table S7) days of the life. The effects of 0.1; 1.0 and 5.0 mg/ml of chokeberry extract were analyzed (see Fig. 7).

**Fig. 7.**
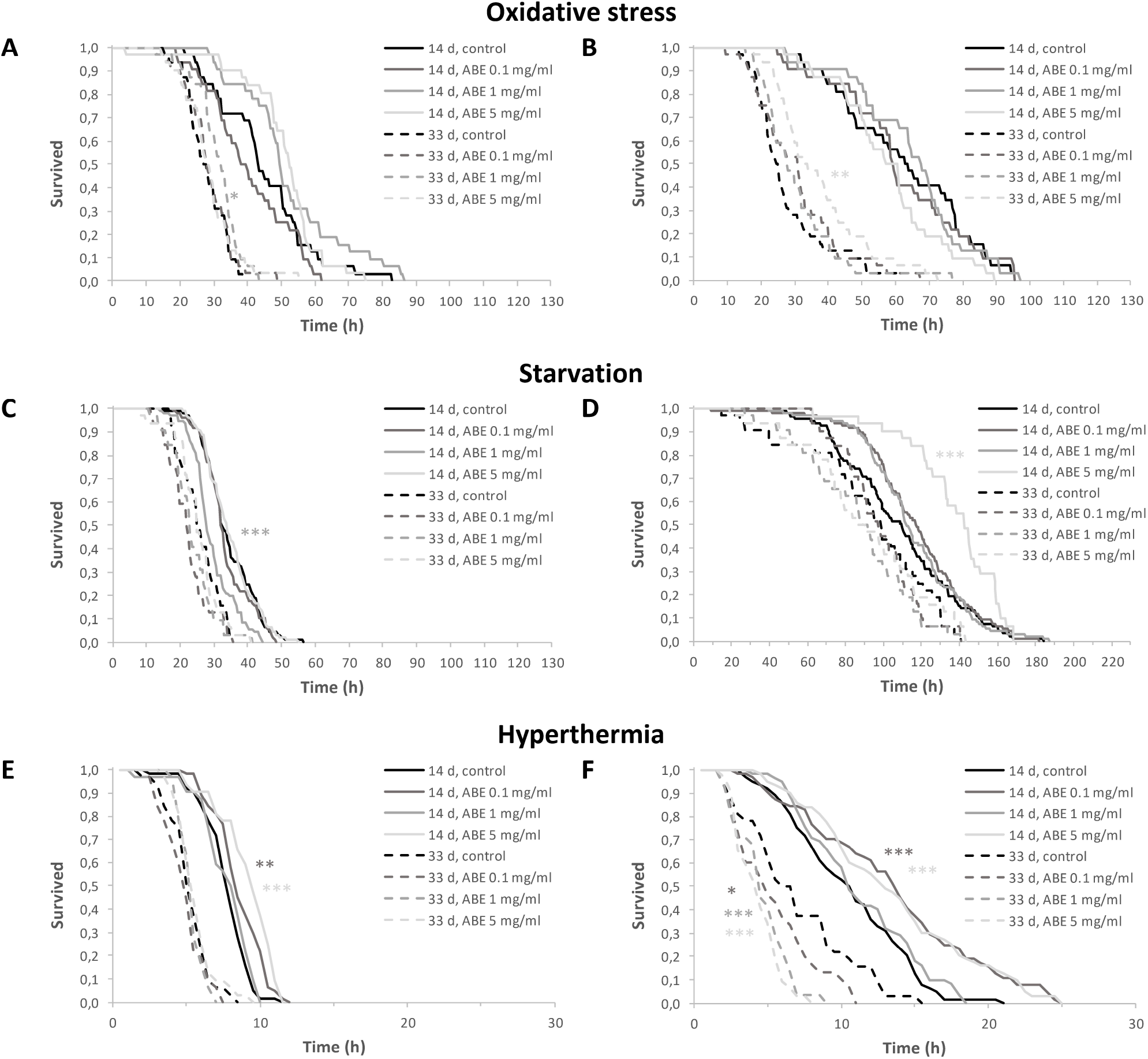
Effects of ABE treatment on the relative expression level of stress response genes in male (A, C) and female (B, D) flies at the age 14 days (A, C) and 33 days (B, D). *p<0.05, **p<0.01, ***p<0.001, t-Student test. All diagrams represent the means of three biological replicates, each quantification is carried out in three technical replicates, the error bars show the standard error of the mean.

It was revealed that in males at the age of 33 days old (Fig. 7C) 0.1 mg/ml of ABE increased the expression of *Clk* gene, while the transcriptional activity of *per* gene decreased. At the same time, females exhibited increased expression of *per* at concentrations of 0.1 and 5.0 mg/ml. And also on the 33rd day of life, there was a slight increase in *Clk* expression level at all ABE concentrations used. At the age of 14 days old, males (Fig. 7A) had a pronounced increase in the expression of oxidative stress resistance genes such as *Keap1* (0.1 mg/ml) and *sirt1* (1.0 mg/ml) and the expression of these genes decreased with age. In females of different ages, the changes in the expression level of the most studied genes were not significant, except for *Keap1*, which demonstrated the increase of the expression level after supplementation with 0.1 mg/ml of ABE on day 14. With age (33 days of life) (see Fig. 7C), an increase in the expression level of *Hsp68* was caused by 0.1. In females, on the 14th day of life (Fig. 7B), an increase in the expression level of the *Hsp83* gene was observed at 0.1 mg/ml of ABE, at the same time, on the 33rd day of life (Fig. 7D), there was more pronounced increase in the expression level of *Hsp27* at 0.1 mg/ml than other concentrations. With age, females show a general increase in the expression level of genes encoding heat shock proteins.

We demonstrated that the Aronia extract increased expression of the *sirt1* gene that previously showed to be involved in cellular stress defense mechanisms such as protection against oxidative stress, DNA repair and may contribute to decrease in risk of various age-related disorders (Elibol and Kilic 2018; Kayashima et al. 2017). The heat shock proteins (HSP) is one of the systems of cellular defense in response to various stress conditions. Increased expression of HSP genes such as *Hsp27, Hsp68* and *Hsp83* may increase resistance to paraquat and starvation (Wang et al. 2004b). It has been shown that Hsps are involved in acquired resistance to stress (hormesis), such hormesis correlates with increased Hsp expression and may lead to a slight increase in lifespan (Tower 2011). Our data on increased expression of heat shock proteins genes also show a connection with these statements. It should be noted that the effects of HSPs overexpression on longevity are ambiguous. For example overexpression of *hsp70* in the nervous system and muscles or RNAi-mediated down-regulation of *hsp70* had shown no effect on *Drosophila* longevity while ubiquitous overexpression reduced the lifespan of males (Xiao et al. 2019).

The treatment of cells with anthocyanin-rich extract of *Aronia melanocarpa* shown to increase the level of Nrf2 in a concentration-dependent manner. The extract increased activation of Nrf2 approximately 4.5 times in a concentration of 25 µg/ml in comparison with angiotensin II treated cells (Parzonko et al. 2015). Treatment of a glioblastoma cell line (U373) with an extract of chokeberry (*Aronia melanocarpa*) and curcumin (*Curcuma longa*), for 48 hours, caused a decrease in the expression of the MMP-2, −14, −16 and −17 genes(Abdullah Thani et al. 2012). The extract of chokeberry (2.9 g/kg of body weight) supplementation with food and further exercise increased the activity of mTORC1 in rats (Makanae et al. 2019). The the positive effect on lifespan has been reported to be associated with increased expression of *SOD*, *CAT*, *GPx* genes and suppression of the *MTH* gene (Jo and Imm 2017; Lashmanova et al. 2019; Peng et al. 2011; Peng et al. 2012; Wang et al. 2015). Antioxidant activity (decrease in the level of Sod2, Gpx1 and Prdx1) of 4.5% lyophilized powder of chokeberry berries was also revealed in the colon and mesenteric lymph node (Pei et al. 2019a; Pei et al. 2019b). Ethanol extract of chokeberry has been shown to inhibit hydrogen peroxide-induced ROS production in murine macrophage cell culture in a dose-dependent manner and exhibit strong radical scavenging activity (Ghosh et al. 2018)

## Discussion

In this study, we found that the extract of chokeberry in small concentrations and for a limited period of time (in our case, two weeks), increases lifespan, which means it corresponds to the idea that chokeberry may be hormetin. In addition, it was found that the black chokeberry extract meets the main criteria of a geroprotector: it is not toxic and does not exhibit toxic properties, it increases the lifespan of the model organism and improves the quality of life. ABE also meets the secondary criteria: it increases resistance to environmental stressors and has prophylactic properties against age-related diseases (Moskalev et al. 2016). Based on this, we can assume that hormetins are geroprotectors.

The interest of society in improving the quality of life and possible longevity is growing every year, and this is possible due to the reduction of age-related diseases (inflammatory processes in the organism) (Jurikova et al. 2017; Kokotkiewicz et al. 2010; Staszowska-Karkut and Materska 2020). Based on the traditional medicine, we can assume that by supplementing our diet with natural products (berries, fruits and vegetables), we can significantly improve the health of our organism with age (Belwal et al. 2017; Kokotkiewicz et al. 2010; Kristo et al. 2016; Prior 2010) Besides, with age dietary needs change, especially which nutrients are important and needed for an aging organism (Institute of Medicine Food 2010). As mentioned earlier, a plant-based diet containing substances such as polyphenols, flavonoids, and anthocyanins may have positive effects on lifespan and also can help with anti-aging diseases. However, the amount and timing of nutrient intake are equally important. The optimal effect of diet on aging and disease is usually associated with a narrow dose range. Excessive intake or deficiency of nutrients can have an adverse effect on the organism’s health because the dose – response relationship is non-linear (Evangelakou et al. 2019; Skorupa et al. 2008). With regard to the period of nutrient intake, in old age the use of many nutrients becomes less efficient, therefore, their demand increases. It becomes necessary to add berries, fruits and vegetables to the diet, which contain large amounts of polyphenols, including anthocyanins. Anthocyanins produce a wide range of health benefits such as weight gain, anti-inflammatory, insulin resistance and regulation of immune function (Belwal et al. 2017; Ding et al. 2018; Prior 2010; Valenza et al. 2018). It is suggested that berries and fruits supplement to the diet may not only help reduce the risk of various diseases, such as cancer, cardiovascular, type 2 diabetes, but also improve the quality and longevity of life. (Basu et al. 2010; Calvano et al. 2019; Kalt et al. 2020; Kristo et al. 2016). Perhaps the anthocyanins that contained in Aronia may act as mild stress and activate protective mechanisms in cells.

The obtained data suggest that the most pronounced geroprotective effect of ABE observed when treatment was conducted at the age of 4-6 weeks relative to the other tested regimes (Institute of Medicine Food 2010). Namely, when ABE is used in middle age (4-6 weeks of life) and for a limited period of time (in our case, 2 weeks), the average life expectancy in males and females increases by 5% (0.1 and 5.0 mg/ml). The maximum lifespan of ABE (0.01; 0.1; 2.5 and 5.0 mg/ml) increases by 3% in females, and in males by 9% (0.1 and 5.0 mg/ml) *Drosophila melanogaster*. Other results are observed in females, perhaps this is due to sexual dimorphism. The study showed that genes are expressed differently depending on gender (Parisi et al. 2004). This is probably because females need more nutrients, due to sexual dimorphism, that is females experience constant energy costs for reproduction through continuous laying eggs (Camus et al. 2017; Wu et al. 2020). 0.1 mg/ml increases resistance to oxidative stress on the 33rd day of life of flies. In addition, 5.0 mg/ml is an effective dose for females with regard to locomotor activity, as well as to increase resistance to oxidative stress at 33 days of life, as well as hyperthermia and starvation at 14 days of age. Hormetic reactions associated with the *Nrf2 / Keap* pathway exhibit the geroprotective effect of ethanol extract of chokeberry berries. The same metabolic pathways and cellular processes are activated by polyphenols (Martel et al. 2019). Anthocyanins found in chokeberry fruits, may reduce ROS levels by activating NRF2 thereby inducing the expression of antioxidant enzymes (Belwal et al. 2017; Ding et al. 2018; Prior 2010; Valenza et al. 2018). The induced increase in the lifespan of *Drosophila* Aronia extract could be due to hormetic reactions.

## Conclusions

According to our data, the ethanol extract of chokeberry in low concentrations and in a short time period manifests itself as hormetin, thereby showing the greatest geroprotective potential, on the life cycle of *Drosophila melanogaster*. In addition, it can be assumed that we have found the critical age for dietary intervention, when the addition of Aronia extract maximizes the median and maximum life expectancy. However, for such high-profile statements, more experiments are needed with different extracts and their reproducible results.

But, our result shows that black chokeberry extract contains various antioxidants that may positively affect the longevity of life. In addition, Aronia extract may increase the resistance of flies to hyperthermia and oxidative stress. Aronia extract may become one of the most dignified “weapons” in the fight against aging.

## Author Contributions

Conceptualization, ЕYP, MVS, AАМ; Methodology, ЕYP, NVZ, MVS, VVP, SAP; Software, ЕYP, NVZ, MVS; Investigation, ЕYP, NVZ, MVS, DAG, DVK, NRM, NSU, VVP, SAP; Data Curation, ЕYP, NVZ, MVS; Writing – Original Draft Preparation, ЕYP, NVZ, MVS, DAG; Writing – Review & Editing, AАМ; Visualization, ЕYP, NVZ, MVS, SAP; Supervision, MVS, AАМ; Project Administration, ЕYP, AАМ; Funding Acquisition, AАМ.

## Acknowledgments

We are grateful to the Institute of Chemistry of Komi Science Center for assistance in the analysis of extract composition.

## Compliance with ethical standards

## Electronic supplementary material

Supplementary material 1 (PDF 257 kb)

