## Supplementary Tables for "Geroprotective effects of *Aronia melanocarpa* fruit extract on *Drosophila melanogaster*"

### Electronic Supplementary Materials

#### Supplementary Tables

**Supplementary Table S1** Influence of ethanol extract on lifespan, when the extract is added throughout life of an imago

| Variant, mg/ml | Sex | M, day | dM % | Mantel - Cox test | Wilcoxon-Breslow-Gehan test | 90%, day | d90 % | Wang - Allison test | N |
|---|---|---|---|---|---|---|---|---|---|
| Control | ♂ | 59 | | | | 70 | | | 743 |
| 0.01 | ♂ | 59 | 0 | $p < 0.05$ | $p > 0.05$ | 68 | -2.9 | $p < 0.0001$ | 363 |
| 0.1 | ♂ | 57 | -4.2 | $p < 0.0001$ | $p < 0.0001$ | 64 | -8.6 | $p < 0.0001$ | 378 |
| 1.0 | ♂ | 58 | -1.7 | $p < 0.01$ | $p > 0.05$ | 65 | -7.1 | $p < 0.0001$ | 418 |
| 2.5 | ♂ | 58 | -1.7 | $p < 0.0001$ | $p < 0.001$ | 65 | -7.1 | $p < 0.0001$ | 409 |
| 5.0 | ♂ | 58 | -1.7 | $p < 0.0001$ | $p < 0.01$ | 66 | -5.7 | $p < 0.001$ | 415 |
| 10.0 | ♂ | 58 | -1.7 | $p > 0.05$ | $p > 0.05$ | 72 | 2.9 | $p < 0.0001$ | 371 |
| Control | ♀ | 66 | | | | 76 | | | 741 |
| 0.01 | ♀ | 66 | 0 | $p > 0.05$ | $p > 0.05$ | 76 | 0 | $p > 0.05$ | 349 |
| 0.1 | ♀ | 64 | -3.0 | $p > 0.05$ | $p > 0.05$ | 76 | 0 | $p > 0.05$ | 357 |
| 1.0 | ♀ | 66 | 0 | $p > 0.05$ | $p > 0.05$ | 77 | 1.3 | $p > 0.05$ | 432 |
| 2.5 | ♀ | 64 | -3.0 | $p < 0.0001$ | $p < 0.0001$ | 72 | -5.3 | $p < 0.01$ | 400 |
| 5.0 | ♀ | 65 | -2.3 | $p < 0.01$ | $p < 0.01$ | 75 | -1.3 | $p > 0.05$ | 398 |
| 10.0 | ♀ | 64 | -3.0 | $p > 0.05$ | $p > 0.05$ | 75 | -1.3 | $p > 0.05$ | 375 |

M - median lifespan; 90% - age of 90% mortality (maximum lifespan), dM - difference median lifespan, d90% -difference age of 90% mortality, n - number of flies, ♂ – males, ♀ – females.

**Supplementary Table S2** Effect of ethanol ABE supplementation on lifespan, 1 – 2 weeks old

| Variant, mg/ml | Sex | M, day | dM % | Mantel - Cox test | Wilcoxon-Breslow-Gehan test | 90%, day | d90 % | Wang - Allison test | N |
|---|---|---|---|---|---|---|---|---|---|
| Control | ♂ | 58 | | | | 65 | | | 453 |
| 0.01 | ♂ | 58 | 0 | $p > 0.05$ | $p > 0.05$ | 67 | 3.1 | $p > 0.05$ | 385 |
| 0.1 | ♂ | 58 | 0 | $p > 0.05$ | $p > 0.05$ | 67 | 3.1 | $p > 0.05$ | 407 |
| 1.0 | ♂ | 58 | 0 | $p < 0.05$ | $p > 0.05$ | 68 | 4.6 | $p > 0.05$ | 395 |
| 2.5 | ♂ | 60 | 3.4 | $p < 0.0001$ | $p < 0.0001$ | 68 | 4.6 | $p > 0.05$ | 411 |
| 5.0 | ♂ | 57 | -1.7 | $p > 0.05$ | $p > 0.05$ | 67 | 3.1 | $p > 0.05$ | 386 |
| 10.0 | ♂ | 57 | -1.7 | $p < 0.05$ | $p > 0.05$ | 64 | -1.5 | $p > 0.05$ | 253 |
| Control | ♀ | 66 | | | | 75 | | | 461 |
| 0.01 | ♀ | 64 | -3 | $p < 0.0001$ | $p < 0.001$ | 73 | -2.7 | $p < 0.0001$ | 393 |
| 0.1 | ♀ | 64 | -3 | $p < 0.01$ | $p < 0.001$ | 75 | 0 | $p > 0.05$ | 391 |
| 1.0 | ♀ | 64 | -3 | $p < 0.01$ | $p < 0.0001$ | 75 | 0 | $p > 0.05$ | 386 |
| 2.5 | ♀ | 64 | 0 | $p > 0.05$ | $p > 0.05$ | 76 | 1.3 | $p > 0.05$ | 384 |
| 5.0 | ♀ | 65 | -1.5 | $p < 0.05$ | $p < 0.05$ | 75 | 0 | $p < 0.05$ | 394 |
| 10.0 | ♀ | 60 | -9.1 | $p < 0.0001$ | $p < 0.0001$ | 71 | -5.3 | $p < 0.0001$ | 259 |

M - median lifespan; 90% - age of 90% mortality (maximum lifespan), dM - difference median lifespan, d90% -difference age of 90% mortality, n - number of flies, ♂ – males, ♀ – females.

**Supplementary Table S3** Effect of ethanol ABE supplementation on lifespan, 4 – 6 weeks old

| Variant, mg/ml | Sex | M, day | dM % | Mantel - Cox test | Wilcoxon-Breslow-Gehan test | 90%, day | d90 % | Wang - Allison test | N |
|---|---|---|---|---|---|---|---|---|---|
| Control | ♂ | 56 | 5.4 | | | 65 | | | 461 |
| 0.01 | ♂ | 59 | 5.4 | $p < 0.01$ | $p < 0.01$ | 68 | 4.6 | $p < 0.05$ | 379 |
| 0.1 | ♂ | 59 | 5.4 | $p < 0.0001$ | $p < 0.0001$ | 71 | 9.2 | $p < 0.0001$ | 390 |
| 1.0 | ♂ | 59 | 5.4 | $p < 0.001$ | $p < 0.01$ | 68 | 4.6 | $p < 0.05$ | 375 |
| 2.5 | ♂ | 59 | 5.4 | $p < 0.01$ | $p < 0.01$ | 67 | 3.1 | $p < 0.05$ | 379 |
| 5.0 | ♂ | 59 | 5.4 | $p < 0.0001$ | $p < 0.0001$ | 71 | 9.2 | $p < 0.0001$ | 381 |
| 10.0 | ♂ | 58 | 3.6 | $p > 0.05$ | $p > 0.05$ | 68 | 4.6 | $p > 0.05$ | 237 |
| Control | ♀ | 67 | | | | 76 | | | 460 |
| 0.01 | ♀ | 67 | 0 | $p > 0.05$ | $p > 0.05$ | 78 | 2.6 | $p < 0.0001$ | 383 |
| 0.1 | ♀ | 70 | 4.5 | $p < 0.01$ | $p < 0.05$ | 78 | 2.6 | $p < 0.0001$ | 383 |
| 1.0 | ♀ | 67 | 0 | $p > 0.05$ | $p > 0.05$ | 78 | 2.6 | $p < 0.01$ | 384 |
| 2.5 | ♀ | 68 | 1.5 | $p < 0.01$ | $p > 0.05$ | 78 | 2.6 | $p < 0.0001$ | 376 |
| 5.0 | ♀ | 70 | 4.5 | $p < 0.01$ | $p < 0.05$ | 78 | 2.6 | $p < 0.0001$ | 377 |
| 10.0 | ♀ | 67 | 0 | $p > 0.05$ | $p > 0.05$ | 74 | -2.6 | $p > 0.05$ | 246 |

M - median lifespan; 90% - age of 90% mortality (maximum lifespan), dM - difference median lifespan, d90% -difference age of 90% mortality, n - number of flies, ♂ – males, ♀ – females.

**Supplementary Table S4** The result of the effect of ABE on the number of males and females with the Smurf phenotype at the age of 2, 6 and 8 weeks old

| Male 2 week old | | | | | | Female 2 week old | | | | | |
|---|---|---|---|---|---|---|---|---|---|---|---|
| Variant, mg/ml | normal "-" | smurf "+" | % smurf | error of % sm | Fisher's exact test (p) | Variant, mg/ml | normal "-" | smurf "+" | % smurf | error of % sm | Fisher's exact test (p) |
| control | 118 | 0 | 0 | 0 | n/a | control | 118 | 0 | 0 | 0 | n/a |
| 0.01 | 117 | 0 | 0 | 0 | n/a | 0.01 | 119 | 0 | 0 | 0 | n/a |
| 0.1 | 119 | 0 | 0 | 0 | n/a | 0.1 | 118 | 0 | 0 | 0 | n/a |
| 1.0 | 116 | 0 | 0 | 0 | n/a | 1.0 | 119 | 0 | 0 | 0 | n/a |
| 2.5 | 123 | 0 | 0 | 0 | n/a | 2.5 | 118 | 0 | 0 | 0 | n/a |
| 5.0 | 120 | 0 | 0 | 0 | n/a | 5.0 | 117 | 0 | 0 | 0 | n/a |
| 10.0 | 116 | 0 | 0 | 0 | n/a | 10.0 | 115 | 0 | 0 | 0 | n/a |
| Male 6 week old | | | | | | Female 6 week old | | | | | |
| Variant, mg/ml | normal "-" | smurf "+" | % smurf | error of % sm | Fisher's exact test (p) | Variant, mg/ml | normal "-" | smurf "+" | % smurf | error of % sm | Fisher's exact test (p) |
| control | 143 | 0 | 0 | 0 | n/a | control | 148 | 4 | 2.7 | 1.3 | n/a |
| 0.01 | 146 | 0 | 0 | 0 | n/a | 0.01 | 138 | 4 | 2.9 | 1.4 | 0.1 |
| 0.1 | 132 | 0 | 0 | 0 | n/a | 0.1 | 147 | 0 | 0 | 0.0 | n/a |
| 1.0 | 147 | 0 | 0 | 0 | n/a | 1.0 | 161 | 2 | 1.2 | 0.9 | 0.9 |
| 2.5 | 142 | 1 | 0.7 | 0.7 | 1.4 | 2.5 | 161 | 2 | 1.2 | 0.9 | 0.9 |
| 5.0 | 156 | 0 | 0 | 0 | n/a | 5.0 | 155 | 5 | 3.2 | 1.4 | 0.3 |
| 10.0 | 163 | 0 | 0 | 0 | n/a | 10.0 | 156 | 1 | 0.6 | 0.6 | 1.5 |
| Male 8 week old | | | | | | Female 8 week old | | | | | |
| Variant, mg/ml | normal "-" | smurf "+" | % smurf | error of % sm | Fisher's exact test (p) | Variant, mg/ml | normal "-" | smurf "+" | % smurf | error of % sm | Fisher's exact test (p) |
| control | 87 | 2 | 2.3 | 1.6 | n/a | control | 100 | 3 | 3.0 | 1.7 | n/a |
| 0.01 | 77 | 0 | 0 | 0 | n/a | 0.01 | 104 | 0 | 0 | 0 | n/a |
| 0.1 | 82 | 0 | 0 | 0 | n/a | 0.1 | 116 | 3 | 2.6 | 1.5 | 0.2 |
| 1.0 | 82 | 0 | 0 | 0 | n/a | 1.0 | 111 | 1 | 0.9 | 0.9 | 1.1 |
| 2.5 | 90 | 0 | 0 | 0 | n/a | 2.5 | 83 | 2 | 2.4 | 1.7 | 0.2 |

| | | | | | | | | | | | |
|---|---|---|---|---|---|---|---|---|---|---|---|
| 5.0 | 80 | 3 | 3.8 | 2.1 | 0.6 | 5.0 | 111 | 2 | 1.8 | 1.3 | 0.6 |
| 10.0 | 77 | 0 | 0 | 0 | n/a | 10.0 | 114 | 3 | 2.6 | 1.5 | 0.2 |

*normal* "-" - number of unpainted flies, *smurf* "+" - number of painted flies

*p> 0.05, Fisher's exact test

**Supplementary Table S5** List of primers for qRT-PCR

| Gene | Symbol (FlyBase) | Forward/Reverse (5'-3') |
|---|---|---|
| *eukaryotic translation elongation factor 1 alpha 2* | *eEF1α2* | AGGGCAAGAAGTAGCTGGTTTGC/ GCTGCTACTACTGCGTGTTGTTG |
| *β-Tubulin at 56D* | *Tubulin* | GCAACTCCACTGCCATCC/ CCTGCTCCTCCTCGAACT |
| *Ribosomal protein L32* | *RpL32* | GAAGCGCACCAAGCACTTCATC/ CGCCATTTGTGCGACAGCTTAG |
| *clock circadian regulator* | *clock/Clk* | ATGATGACGCACGTCAGTTCGC/ TCGATGGTGTTCTCGGTGATGC |
| *period* | *per* | GGGATCATATCGCACGTGGAC/ CTGCGGCCAATCAGGTCCTG |
| *Hypoxia-inducible factor 1* | *HIF1/tango/CG1 1987* | TGAGCACAGGCGACCCAAATTAC/ TGTCCTGTATGTTCGCCTCGTC |
| *Kelch-like ECH associated protein 1* | *Keap1* | GCCAATTGGATCCCACGAATGC/ ATTCCTCCTCTTGGCACACCTG |
| *cap-n-collar* | *cnc/NRF2/CG432 86* | GCCAATTGGATCCCACGAATGC/ ATTCCTCCTCTTGGCACACCTG |
| *Superoxide dismutase 1* | *Sod1* | TGCACGAGTTCGGTGACAACAC/ TCCTTGCCATACGGATTGAAGTGC |
| *Heat shock protein 27* | *Hsp27* | ACTGGGTCGTCGTCGTTATTCG/ CGCGCGACGTGACATTTGATTG |
| *Heat shock protein 68* | *Hsp68* | TGGGCACATTCGATCTCACTGG/ TAACGTCGATCTTGGGCACTCC |
| *Heat shock protein 83* | *Hsp83* | AAGATGCCAGAAGAAGCAGAGAC C/ATCTTGTCCAGGGCATCGGAAG |
| *Sirtuin 1 (Silent Information Regulator proteins)* | *Sirt1* | TCCAGGACAGTTAGCAGCAGTG/ GGCTACGATTTCGCAGCTTCTC |

**Supplementary Table S6** Effects of ABE treatment on the relative expression level of stress response genes, 14 days

| Gene | Sex | Control | 0.1 mg/ml | 1.0 mg/ml | 5.0 mg/ml |
|---|---|---|---|---|---|
| *Clk* | ♂ | 1.00±0.16 | 0.52±0.04* | 1.69±0.27* | 0.40±0.09* |
| *per* | ♂ | 1.00±0.15 | 0.28±0.20 | 0.41±0.36 | 0.17±0.06* |
| *Keap1* | ♂ | 1.00±0.18 | 8.15±0.42* | 0.56±1.13* | 0.21±0.27* |
| *NRF2* | ♂ | 1.00±0.19 | 0.40±0.14* | 0.61±0.07* | 0.51±0.10 |
| *HIF1* | ♂ | 1.00±0.27 | 0.44±0.04* | 0.41±0.03 | 0.44±0.19* |
| *Sod1* | ♂ | 1.00±0.15 | 0.98±0.05* | 1.27±0.10 | 0.65±0.03* |
| *Sirt1* | ♂ | 1.00±0.15 | 1.88±0.14* | 8.11±0.37 | 1.34±0.12* |
| *Hsp27* | ♂ | 1.00±0.20 | 0.25±0.07* | 0.62±0.44 | 0.42±0.06* |
| *Hsp68* | ♂ | 1.00±0.21 | 0.13±0.09* | 5.65±0.42* | 0.91±0.07* |
| *Hsp83* | ♂ | 1.00±0.22 | 0.22±0.08 | 0.38±0.16 | 0.24±0.03* |
| *Clk* | ♀ | 1.00±0.07 | 0.29±0.76 | 0.50±0.23 | 0.39±0.04* |
| *per* | ♀ | 1.00±0.07 | 10.47±1.91* | 1.11±0.13 | 4.83±0.75* |
| *Keap1* | ♀ | 1.00±0.09 | 6.57±0.95* | 1.46±0.33 | 1.43±0.45 |
| *NRF2* | ♀ | 1.00±0.14 | 1.06±0.53 | 0.68±0.15* | 0.47±0.13* |
| *HIF1* | ♀ | 1.00±0.10 | 0.68±0.34 | 0.86±0.17 | 1.70±0.20* |
| *Sod1* | ♀ | 1.00±0.05 | 1.33±0.72 | 1.41±0.36 | 0.60±0.06* |
| *Sirt1* | ♀ | 1.00±0.06 | 1.42±0.41 | 1.43±0.31 | 1.49±0.16* |
| *Hsp27* | ♀ | 1.00±0.07 | 1.22±0.10* | 2.05±0.27* | 0.72±0.09* |

| | | | | | |
|---|---|---|---|---|---|
| *Hsp68* | ♀ | 1.00±0.15 | 0.78±0.38 | 0.20±0.09* | 0.10±0.02* |
| *Hsp83* | ♀ | 1.00±0.06 | 4.81±0.62* | 0.69±0.12* | 0.29±0.02* |

*p<0.05, t-Student test. Errors indicate standard error of the mean. ♂ – males, ♀ – females.

**Supplementary Table S7** Effects of treatment on the relative expression level of stress response genes 33 days

| Gene | Sex | Control | 0.1 mg/ml | 1.0 mg/ml | 5.0 mg/ml |
|---|---|---|---|---|---|
| *Clk* | ♂ | 1.00±0.14 | 1.53±0.17* | 0.73±0.08 | 0.60±0.05* |
| *per* | ♂ | 1.00±0.09 | 0.17±0.02* | 0.23±0.04* | 0.32±0.04* |
| *Keap1* | ♂ | 1.00±0.16 | 0.88±0.23* | 0.28±0.04* | 0.07±0.01* |
| *NRF2* | ♂ | 1.00±0.12 | 0.62±0.09* | 1.12±0.16* | 1.38±0.09 |
| *HIF1* | ♂ | 1.00±0.12 | 0.33±0.06* | 0.44±0.04* | 0.26±0.02* |
| *Sod1* | ♂ | 1.00±0.06 | 1.10±0.03 | 1.11±0.08* | 1.22±0.05* |
| *Sirt1* | ♂ | 1.00±0.08 | 0.31±0.24* | 0.61±0.06 | 0.61±0.03* |
| *Hsp27* | ♂ | 1.00±0.11 | 0.31±0.03* | 0.50±0.05* | 0.32±0.01* |
| *Hsp68* | ♂ | 1.00±0.17 | 4.19±0.17* | 3.48±0.34* | 1.25±0.07* |
| *Hsp83* | ♂ | 1.00±0.07 | 1.25±0.08 | 1.35±0.06 | 1.32±0.05* |
| *Clk* | ♀ | 1.00±0.09 | 1.17±0.11 | 1.09±0.11 | 1.45±0.17* |
| *per* | ♀ | 1.00±0.13 | 0.84±0.03* | 1.78±0.23 | 2.26±0.25* |
| *Keap1* | ♀ | 1.00±0.25 | 2.20±0.37* | 2.08±0.30* | 1.64±0.24* |
| *NRF2* | ♀ | 1.00±0.15 | 1.58±0.11* | 1.24±0.18 | 1.75±0.25* |
| *HIF1* | ♀ | 1.00±0.13 | 1.80±0.25* | 1.07±0.13* | 1.65±0.18* |
| *Sod1* | ♀ | 1.00±0.15 | 1.60±0.11* | 1.26±0.16* | 1.54±0.15* |
| *Sirt1* | ♀ | 1.00±0.11 | 1.08±0.07* | 0.86±0.13 | 1.13±0.11* |

| | | | | | |
|---|---|---|---|---|---|
| *Hsp27* | ♀ | 1.00±0.14 | 3.45±0.18 | 2.64±0.15* | 2.26±0.21* |
| *Hsp68* | ♀ | 1.00±0.12 | 2.24±0.28 | 2.07±0.70 | 1.63±0.18 |
| *Hsp83* | ♀ | 1.00±0.09 | 1.51±0.09* | 1.34±0.16 | 1.81±0.23* |

*$p<0.05$, t-Student test. Errors indicate standard error of the mean. ♂ – males, ♀ – females
